# ABHD4 regulates adipocyte differentiation in vitro but does not affect adipose tissue lipid metabolism in mice

**DOI:** 10.1101/2022.11.29.518076

**Authors:** Mary Seramur, Sandy Sink, Anderson O’Brien Cox, Cristina M. Furdui, Chia-Chi Chuang Key

## Abstract

Alpha/beta hydrolase domain-containing protein 4 (ABHD4) catalyzes the deacylation of *N*-acyl phosphatidyl-ethanolamine (NAPE) and Lyso-NAPE to produce Glycerophospho-*N*-acyl ethanolamine (GP-NAE). Through a variety of metabolic enzymes, NAPE, Lyso-NAPE, and GP-NAE are ultimately converted into NAE, a group of bioactive lipids that control many physiological processes including inflammation, cognition, and food intake (i.e., oleoylethanolamide or OEA). In a diet-induced obese mouse model, adipose ABHD4 gene expression positively correlated with adiposity. However, it is unknown whether ABHD4 is a causal or a reactive gene to obesity. To fill this knowledge gap, we generated an ABHD4 knockout (KO) 3T3-L1 pre-adipocyte. During adipogenic stimulation, ABHD4 KO pre-adipocytes had increased adipogenesis and lipid accumulation, suggesting ABHD4 is responding to (a reactive gene), not contributing to (not a causal gene), adiposity and may serve as a mechanism for protecting against obesity. However, we did not observe any differences in adiposity and metabolic outcomes between adipocyte specific ABHD4 KO mice or whole body ABHD4 KO mice and their littermate control mice on chow or a high fat diet. This might be because we found that deletion of ABHD4 did not affect NAE such as OEA production, even though ABHD4 was highly expressed in adipose tissue and correlated with fasting adipose OEA levels and lipolysis. These data suggest that ABHD4 plays a role in adipocyte differentiation in vitro but not in adipose tissue lipid metabolism in mice despite nutrient overload, possibly due to compensation from other NAPE and NAE metabolic enzymes.

## Introduction

In the United States, from 1999 through 2018, the prevalence of adult obesity increased from 30.5% to 42.4%(1). The most common chronic conditions associated with adult obesity include type 2 diabetes, cardiovascular disease, stroke, and cancer which overall cause about 1 in 5 deaths in the United State each year(2, 3). Obesity is caused by a chronic imbalance between energy intake and energy expenditure resulting in an increase in both adipocyte number (i.e., hyperplasia) and adipocyte size (i.e., hypertrophy)(4, 5). The worldwide epidemic of obesity has spurred increased interest in understanding the molecular mechanisms regulating adipocyte hyperplasia and hypertrophy and in developing new therapeutic approaches(4, 5).

The alpha/beta hydrolase domain (ABHD) family of enzymes plays an important role in the regulation of lipid metabolism and signal transduction(6, 7). For example, ABHD5, also known as Comparative gene identification 58 (CGI-58), is highly expressed in adipose tissue, and during lipolysis, ABHD5/CGI-58 coactivates Adipose triglyceride lipase (ATGL) to catalyze triacylglycerol (TAG) hydrolysis, resulting in free fatty acid and glycerol release into the circulation. Mutation of ABHD5 disrupts lipolysis leading to adiposity and ectopic lipid accumulation in *Arabidopsis*(8), C. elegans(9), mice(10), and humans (i.e., Chanarin-Dorfman syndrome)(11).

ABHD4 is a paralog of ABHD5 (ABHD4 shares 50-55% sequence identity with ABHD5), but it does not affect ATGL activity. In 2006, Simon and Cravatt first identified that ABDH4 is a Lyso-phospholipase/phospholipase B that catalyzes the deacylation of *N*-acyl phosphatidylethanolamine (NAPE) and Lyso-NAPE to produce Glycerophospho-*N*-acyl ethanolamine (GP-NAE)(12), an alternative pathway compared to a classical NAPE-phospholipase D (NAPE-PLD) pathway (**Fig. 1A**). They further described that Glycerol-phosphodiester phosphodiesterase 1 (GDE1) catalyzes the hydrolysis of GP-NAE to produce glycerol-3-phosphate and NAE(13) (**Fig. 1A**). NAE is a group of bioactive lipids that regulate many physiological processes including pain, inflammation, anxiety, cognition, and food intake(14). Recently, the Ueda lab revealed two new enzymes, GDE4 (also known as Glycerophosphodiester phosphodiesterase domain containing 1 or GDPD1)(15) and GDE7 (also known as GDPD3)(16), that can convert Lyso-NAPE to Lyso-phosphatidic acid (Lyso-PA) and NAE (**Fig. 1A**).

**Fig. 1.**
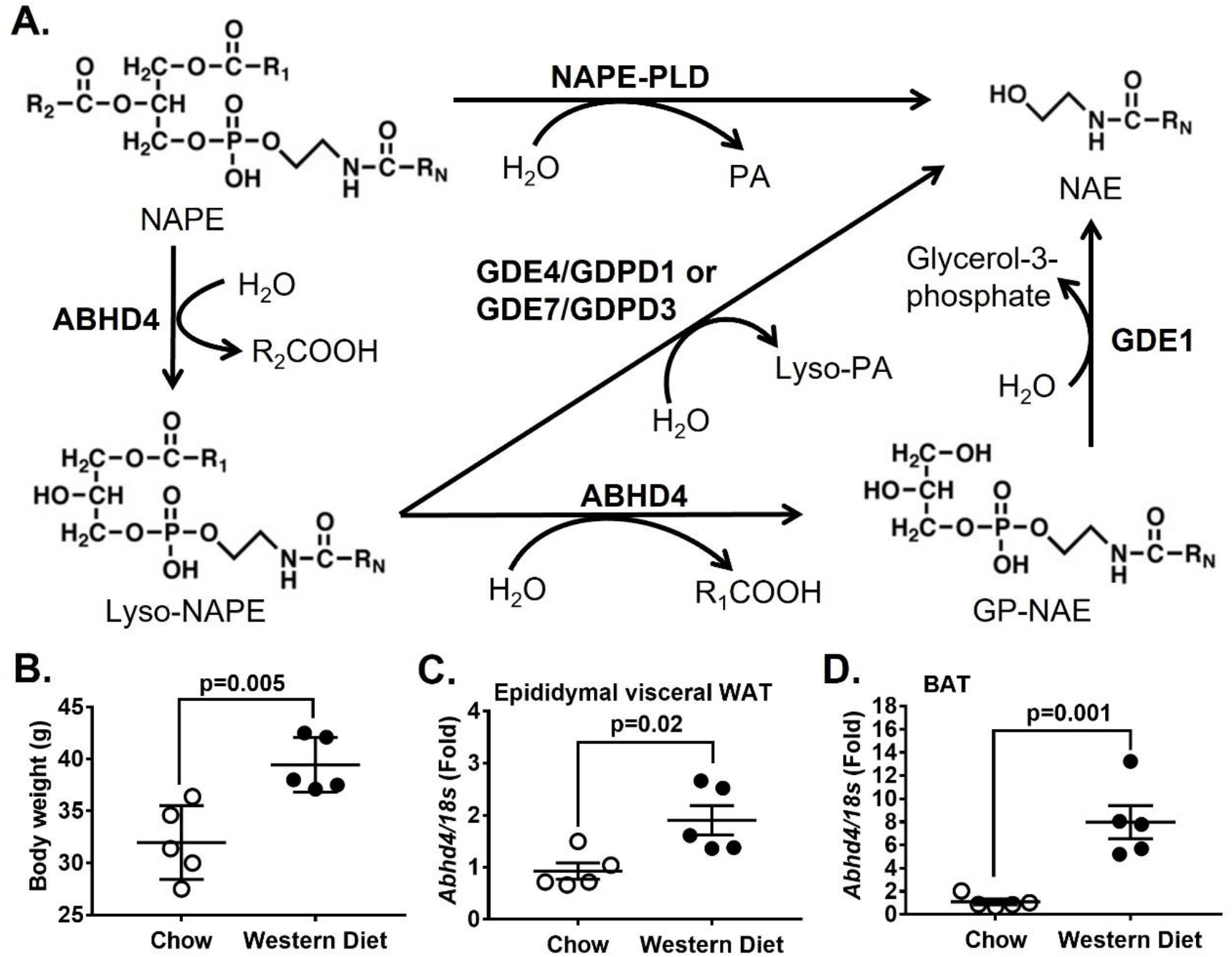
ABHD4’s role in lipid metabolism and its relationship to adiposity. **(A)** The classical pathway to convert *N*-acyl phosphatidyl-ethanolamine (NAPE) to *N*-acyl ethanolamine (NAE) is through NAPE-phospholipase D (NAPE-PLD). Enzymes involved in an alternative pathway or NAPE-PLD-independent pathway of NAE production, including Alpha/beta hydrolase domain-containing protein 4 (ABHD4), Glycerol-phosphodiester phosphodiesterase 1 (GDE1), GDE4/Glycerophosphodiester phosphodiesterase domain-containing protein 1 (GDPD1), and GDE7/GDPD3. ABHD4 catalyzes the deacylation of NAPE and Lyso-NAPE to produce Glycerophospho-*N*-acyl ethanolamine (GP-NAE). GDE1 catalyzes the hydrolysis of GP-NAE to produce glycerol-3-phosphate and NAE. GDE4/GDPD1 and GDE7/GDPD3 can convert Lyso-NAPE to Lyso-phosphatidic acid (Lyso-PA) and NAE. **(B)** Eight-week-old male C57BL/6J mice were fed chow or a Western diet for 16 weeks (n=5 per diet group) and their body weight was measured. Mice were then fasted for 24 hours and their **(C)** epididymal white adipose tissue (WAT) and **(D)** interscapular brown adipose tissue (BAT) RNA was extracted and reverse-transcribed into cDNA for real-time PCR quantification of *Abhd4* normalized to *18s* (endogenous control). All results are mean ± SEM, presented as the fold change compared to chow-fed mouse group, and analyzed using a two-tailed Student’s unpaired *t*-test.

Studies have shown that deletion of NAPE-PLD in hepatocytes and adipocytes result in TAG accumulation in the liver (i.e., steatosis)(17) and adipose tissue (i.e., adiposity/obesity)(18), respectively. However, the underlying mechanisms are still unclear. In addition, our group and others have shown that GDE1(19) and GDE7/GDPD3(20) regulate liver TAG metabolism by controlling the production of glycerol-3-phosphate and Lyso-PA, intermediate substrates of the Glycerol Phosphate Pathway for *de novo* TAG synthesis, respectively. A whole body ABHD4 KO mouse created using a standard gene targeting strategy has been studied(21), however, it is still unclear whether ABHD4 is involved in adipose NAPE and NAE metabolism and whether it affects any mouse metabolic phenotypes of mice. Therefore, in this study, we wanted to investigate the functional role of ABHD4 in adipocyte differentiation and adipose tissue lipid metabolism. To do so, we have generated a new ABHD4 knockout (KO) 3T3-L1 pre-adipocyte and a new floxed Abhd4 mouse for the creation of an adipocyte specific ABHD4 KO mouse. We also obtained a new whole body ABHD4 KO mouse created using CRISPR gene editing from the Jackson Laboratory.

## Methods

### Cell culture

The 3T3-L1 pre-adipocyte cell line was purchased from ATCC (CL-173™). The ABHD4 KO 3T3-L1 pre-adipocyte was generated using CRISPR gene editing and provided by Synthego Corporation (cells are available upon request). The guide sequence (i.e., TTATGTATCCCTCCCAAACC) was designed to target exon 3 of Abhd4 (**Supplementary Fig. 1A**). The KO clone was cut and had an extra nucleotide (i.e., T) insertion (**Supplementary Fig. 1A**) during the non-homologous end joining repair process, resulting in a frameshift mutation that causes premature termination of translation at a new nonsense codon, as confirmed by the Sanger sequence.

The wildtype (WT) control and ABHD4 KO 3T3-L1 pre-adipocytes were cultured and differentiated into adipocytes as described previously(22). Briefly, pre-adipocytes were seeded at 0.05 × 10^6^ cells per well in a 6-well culture plate (for RNA extraction, microscopic imaging, and radioisotope experiments), and 0.1 × 10^6^ cells per 60 mm culture dish (for Oil Red O staining), and 0.2 × 10^6^ cells per 100 mm culture dish (for protein collection and TAG mass analysis). Cells were cultured in Dulbecco’s Modified Eagle Medium (DMEM, Gibco) supplemented with 10% iron fortified calf serum (Sigma-Aldrich) and 1% Penicillin/Streptomycin (P/S, Gibco) for 48 hours until ~90% cell confluence. Adipogenesis (Day 0) was induced by changing medium to DMEM containing 10% fetal bovine serum (FBS, Sigma-Aldrich) plus an adipogenic cocktail (Sigma-Aldrich) including 1 μg/ml of insulin, 0.25 μM of dexamethasone, 0.5 mM of 3-isobutyl-1-methylxanthine, and 2 μM of rosiglitazone for 3 days (Day 3). Cells were then treated with 1 μg/ml of insulin only for 3 days (Day 6) and then without any adipogenic reagents for the next 3 days (Day 9). Medium was changed every two days.

### RNA extraction and real-time PCR

Total RNA was harvested from cells and tissues using Qiazol Lysis Reagent and isolated by following the protocol described in the RNeasy Lipid Tissue Mini Kit (Qiagen). Concentration and quality of RNA was determined using a Nanodrop (ThermoFisher Scientific) and standardized to 1 μg of RNA for cDNA synthesis. cDNA was prepared with the OmniScript RT Kit (Qiagen) and stored at −20° C until gene expression analysis using real-time PCR. Real-time PCR was performed in duplicate on the 7500 Real-Time PCR Systems using TaqMan^®^ Fast Advanced Master Mix and TaqMan^®^ gene expression assays (ThermoFisher Scientific) including *Abhd4* (Mm00506368), *Pparg* (Mm0040940_m1), *Srebf1* (Mm00550338_m1), *Acc1* (Mm01304257_m1), *Fasn* (Mm00662319_m1), *Fabp4* (Mm00445878_m1), *Cd36* (Mm00432403_m1), *Agpat9* (Mm04211965), *Agpat6* (Mm04211965), *Agpat1* (Mm0047900_m1), *Agpat2* (Mm00458880_m1), *Dgat1* (Mm00515643_m1), and *Dgat2* (Mm00499536_m1). Gene expression was normalized to the housekeeping gene *18S* rRNA (REF 4352655) and analyzed using the 2^ddCt^ method with 95% confidence (20).

### Lipid extraction and TAG measurement

Total lipids were extracted from WT and ABHD4 KO cells on Day 0, Day 3, Day 6 and Day 9 of adipocyte differentiation. Cells were washed with ice-cold 1X Dulbecco’s phosphate-buffered saline with no added magnesium or calcium (DPBS, Gibco) twice and lipid-extracted with hexane:isopropanol (3:2, vol:vol) overnight at room temperature. The lipid extracts were dried under a nitrogen stream at 60°C. Once dry, 1% Triton-X in chloroform was added and then dried down again. The residue was re-suspended in ddH_2_O and heated at 60° C for 1 hour to yield an aqueous lipid extract for each sample. The TAG levels were measured using an enzymatic assay (Wako Diagnostics L-Type TG M) according to manufacturer instructions. After lipid extraction, cells were dissolved with 0.1 N of NaOH, and protein concentration was measured using a Pierce™ BCA Protein Assay Kit (Life Technologies) for protein normalization (23).

### Microscopic imaging and Oil red O staining

WT and ABHD4 KO cells were imaged on the 6-well plate at 10X magnification with a set scale of 100 μm using Nikon eclipse TE2000 inverted microscope. Another set of cells were used for Oil Red O staining. Briefly, cells were washed twice with 1X DPBS. Cells were then fixed in 10% phosphate-buffered formalin for 30 minutes at room temperature. After fixation, the cells were washed twice again with 1X DPBS and then treated with Oil Red O stain, four parts ddH_2_O with six parts Oil Red O solution (Sigma-Aldrich), for 15 minutes. Each well was gently washed 5 times with ddH_2_O and then imaged (23).

### Fatty acid uptake and lipogenesis

Fatty acid uptake and incorporation into lipids as well as *de novo* lipogenesis were determined using [9,10-^3^H(N)]-oleic acid and [1,2-^14^C]-acetic acid (PerkinElmer), respectively, following the procedure adapted from our previous study(23). Day 7 differentiated WT and ABHD4 KO 3T3-L1 adipocytes were labeled with 0.5 μCi [14C]-acetic acid or 5 μCi [3H]-oleic acid plus 0.04 mM oleic acid (Sigma-Aldrich) conjugated with 0.01 mM fatty acid free-bovine serum albumin (BSA) per ml of DMEM supplemented with 10% FBS, 1% P/S and 100 nM insulin for 0 (no radioisotopes), 30, 60 and 120 minutes. Following radiolabeling, cells were washed with ice-cold DPBS twice and lipid-extracted with hexane:isopropanol (3:2, vol:vol). Lipid classes from standards and cell lipid extracts were separated by thin layer chromatography (TLC) using Silica Gel plates and a solvent system containing hexane:diethyl ether:acetic acid (80:20:2, vol:vol:vol). Lipids were visualized by exposure to iodine vapor, and bands corresponding to TAG, free cholesterol (FC), cholesteryl ester (CE), and phospholipid (PL) were scraped and counted using a scintillation counter. After lipid extraction, cell residue was dissolved with 0.1 N of NaOH, and protein concentrations were measured using a Pierce™ BCA Protein Assay Kit for protein normalization of data.

### Western blot

Cellular protein was harvested in Pierce™ IP lysis buffer (ThermoFisher Scientific) from Day 0 of WT and ABHD4 KO cells. Protein concentration of each sample was determined through the Pierce™ BCA Protein Assay Kit. A total of 20 μg cell protein was separated on a 4-20% polyacrylamide gel (Bio-Rad), then transferred to a nitrocellulose membrane (Bio-Rad). The membrane was blocked using 5% non-fat milk in Tris-buffered saline plus 0.1% Tween (TBST) for 2 hours. Membranes were incubated overnight at 4°C in primary antibodies including ABHD4 (Sigma-Aldrich, SAB1307058) and GAPDH (Santa Cruz Biotechnology, sc-477724). After incubation in primary antibodies, membranes were washed for 15 minutes using TBST and incubated for 1.5 hours in secondary antibodies to produce a chemiluminescent signal. The membranes were washed again and placed in SuperSignal™ West Pico PLUS Chemiluminescent Substrate (Fisher) prior to imaging. Membranes were visualized for proteins of interest using the ChemiDoc Gel Imaging System (Bio-Rad)

### Mice

Mice were housed in standard cages under a 12-hour light cycle and 12-hour dark cycle (dark from 6:00 PM to 6:00 AM) at standard ambient temperature and humidity conditions, and were provided with ad libitum water and a standard chow diet (Purina-LabDiet, Prolab RMH 3000). All experiments were performed using a protocol approved by the Institutional Animal Care and Use Committee at Wake Forest School of Medicine in facilities approved by the American Association for Accreditation of Laboratory Animal Care.

In **Fig. 1B-D**, eight-week-old male C57BL/6J mice (Jackson Laboratory #000664) were fed chow or a Western diet (Envigo #TD 88137, 42% from fat, 0.2% total cholesterol) for 16 weeks (n=5 per diet group). At the time of mouse necropsy, epidydimal visceral white and interscapular brown adipose tissues were harvested and stored in −80°C until used for RNA extraction, cDNA synthesis, and real-time PCR for gene expression. In **Fig. 7A-G**, sixteen-week-old male (n=10) and female (n=10) C57BL/6J mice were fasted for 24 hours (n=5/sex) or fasted for 24 hours and then refed chow for 12 hours (n=5/sex). Male and female gonadal visceral white and interscapular brown adipose tissues were harvested for gene expression using real-time PCR and oleoylethanolamide (OEA) quantification using mass spectrometry. Mouse blood was also collected for quantifying free fatty acid concentrations using an enzymatic colorimetric assay based on the manufactory procedures (Zenbio).

An floxed Abhd4 (Abhd4^flox/flox^) mouse model was generating by flanking exons 3-4 of Abhd4 gene locus and an inverted reporter cassette of 3_SA_IRES_eGFP (3_Splice Acceptor_Internal Ribosome Entry Site_enhanced Green Fluorescent Protein), with loxP and loxP_2272 sites, via gene targeting in C57BL/6 mouse embryonic stem cells (**Supplementary Fig. 1B**) and provided by Ozgene (mice are available upon request). Cre recombinase-mediated deletion of the floxed region should excise exons 3-4 and invert the GFP knockin cassette (3_SA_IRES_eGFP) into the correct orientation for expression (**Supplementary Fig. 1C**). Exons 3 and 4 encode part of the alpha/beta hydrolase fold-1 and loss of this protein domain should eliminate hydrolase activity. In addition, along with deletion of exon 3 and 4, transcription and translation should be terminated with the inclusion of GFP-PolyA cassette, resulting in expression of a truncated protein. Lastly, adipocytes lacking ABHD4 expression can be identified by GFP detection using microscopy.

Abhd4^flox/flox^ mice were crossed with hemizygous Adiponectin-Cre (Adipoq-Cre^+/−^) mice (Jackson Laboratory #028020) (24) for the creation of adipocyte specific ABHD4 KO (Abhd4^flox/flox^Adipoq-Cre^+/−^ as Abhd4^adipose−/−^) and their littermate control (Abhd4^flox/flox^Adipoq-Cre^−/−^ as Abhd4^adipose+/+^). The mouse genotype was determined using genotyping protocols provided by Ozgene (**Table 1** for Abhd4^flox/flox^) and the Jackson Lab (Protocol #20627 for Adipoq-Cre^+/−^) on the 7500 Real-Time PCR Systems.

**Table 1.**
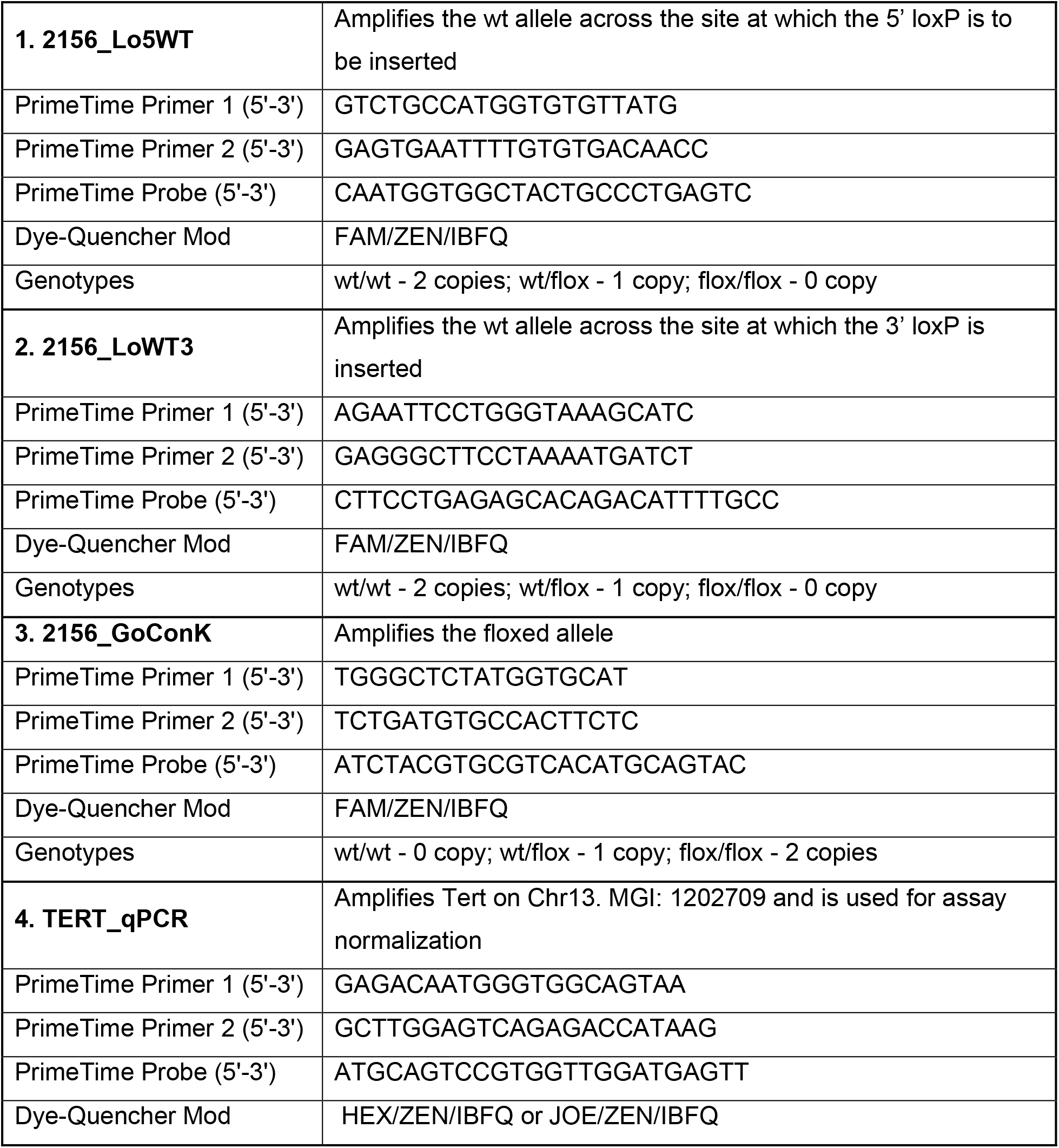
Primers for genotyping a floxed Abhd4 mouse (Integrated DNA Technologies)

In **Fig. 5**, eight-week-old male and female Abhd4^adipose−/−^ and Abhd4^adipose+/+^ mice were maintained on chow or fed a high fat diet (i.e., Research Diets, 45% kcal from fat) for 12 weeks to induce obesity (n=7-9/sex/genotype/diet). Mouse body weight were recorded weekly. Between 8-12 weeks of the experimental diet feeding, mice were used for systemic metabolic phenotyping including 1) glucose and/or insulin tolerance tests (23): Mice were fasted for 16 hours for an intraperitoneal glucose (Sigma-Aldrich) tolerance test (1 g/kg body weight) and fasted for 4 hours for an intraperitoneal insulin (Eli Lilly) tolerance test (0.75 U/kg body weight), and 2) indirect calorimetry (25): Measurement of oxygen consumption, carbon dioxide production, respiratory exchange ratio, energy expenditure, food/water intake, and activity of mice during a 12-hour light/12-hour dark cycle for 5 consecutive days using a TSE PhenoMaster system. Prior to necropsy, mice were fasted overnight, and blood was collected to measure serum insulin levels. Brown and white adipose tissues such as inguinal subcutaneous and gonadal visceral fat pads from male and female mice were collected and weighted. Adipose tissues were snap frozen in liquid nitrogen and stored at −80°C until used for gene and protein expression by real-time PCR and Western blot, respectively.

Another portion of white epidydimal visceral fat pads was used for adipose tissue digestion(26). Briefly, tissue was enzymatically digested in digestion buffer (0.5 g of fat in 10 ml) including 0.8 mg/ml of collagenase II (Worthington Biochemicals), 3% of BSA (Sigma-Aldrich), 1.2 mM of calcium chloride (Sigma-Aldrich), 1 mM of magnesium chloride (Sigma-Aldrich), and 0.8 mM of zinc Chloride (Sigma-Aldrich) in Hanks Buffered Salt Solution (Life Technologies) for 60 minutes at 37° C in a shaking water bath with ~200 rpm agitation. The fat digest was then filtered through a 200-um filter. The stromal vascular fraction (SVF) was pelleted by centrifugation at 800×g for 10 minutes. Red blood cells were lysed using ACK lysis buffer. SV cells were resuspended in 1 ml of medium prior to proliferation and differentiation into adipocytes as described above in Cell Culture section.

In **Fig. 6**, heterozygous Abhd4^+/−^ mice (#46224-JAX) were purchased from the Jackson Laboratory and crossed to generate whole body ABHD4 KO (Abhd4^−/−^) and their WT (Abhd4^+/+^) littermate control mice. Mouse genotype was determined using a genotyping protocol (Protocol 34917) provided by the Jackson Laboratory. Six-week-old male (n=10-13/genotype) and female (n=9-10/genotype) Abhd4^−/−^ and Abhd4^+/+^ mice were fed a high fat diet (i.e., Research Diets, 45% kcal from fat) for 19-24 weeks to induce obesity. Mouse body weight were recorded weekly. Serial measurements of mouse body composition (i.e., whole body fat, lean, free water, and total water masses in live animals) was obtained using EchoMRI™. We then performed systemic metabolic phenotyping and mouse necropsy on these mice as described above.

### Mass spectrometry

Tissues weighed out into a homogenization tube with 2.8 mm ceramic beads before the addition of 1 ml of methanol and 10 μl of internal standard solution (200 pg/μl OEA-d4). The samples were then homogenized using a Bead Ruptor 24 (OMNI International) for 3 cycles of 20 seconds. These homogenized samples were then centrifuged for 10 minutes at 16,000 × g. Supernatants were transferred into glass vials and diluted with 4 ml of 0.1% formic acid, then mixed thoroughly.

Reverse-phase cartridges (Waters Sep-Pak Vac 1cc (100mg) tC18) were activated with 1 ml of methanol and then equilibrated with 3 ml of 0.1% formic acid. Samples were then loaded onto the columns. Columns were rinsed twice, first with 1 ml of 0.1% formic acid, then with 1 ml of 15% methanol. Samples were finally eluted with 2 steps of methanol in 500 μl aliquots.

OEA was quantified using either a Thermo Q Exactive HF hybrid quadrupole-Orbitrap mass spectrometer or a Shimadzu UHPLC-MS/MS. The Q Exactive HF was equipped with a heated electrospray ion (HESI) source and a Vanquish UHPLC System. The Shimadzu instrument was equipped with Nexera UHPLC system and an 8050 triple-quadrupole mass spectrometer utilizing a DUIS source.

Chromatographic methods were identical for both instruments and the compounds were separated on either instrument with a Kinetex C8 (Phenomenex, 150 × 3 mm, 2.6 μm) with a flow rate of 0.4 ml/minute and mobile phases consisting of 0.1% formic acid for mobile phase A and acetonitrile for mobile phase B. The mobile phase gradient began at 30% B and was then increased to 95% B at 9 minutes. This percentage was held for 2 minutes before being decreased to 30% B at 11.1 minutes and then held at that final percentage until 15 minutes.

Ionization occurred at the DUIS source set to the following parameters: nebulizing gas flow of 2 L/min, heating gas flow of 10 L/min, interface temperature of 300°C, DL temperature of 250°C, heat block temperature of 400°C, and a drying gas flow of 10 L/min. Each analyte and its deuterated internal standard used unique MRM transitions in positive ESI mode: OEA 330.20>66.10 and OEA-d4 326.20>62.10.

### Statistics

Data were presented as mean ± standard error of the mean (SEM) throughout the figures. All data points reflected biological replicates. Binary comparisons were performed using two-tailed Student’s t-tests. Datasets comparing the effect of a single independent variable on more than two groups were assessed by one-way ANOVA followed by Dunnett’s correction. Datasets containing groups defined by two independent variables (genotype, time) were assessed by two-way ANOVA with Sidak’s correction. Prism 7 software (GraphPad) is used to perform statistical analyses (Statistical significance p<0.05) and generate graphical representations of data. CalR, a web-based tool, was used to perform statistical analyses (Statistical significance p<0.05) and generate graphical representations of indirect calorimetry data(27).

## Results

### The mRNA levels of Abhd4 in adipose tissue is positively associated with obesity

We first examined gene expression of NAPE-PLD, ABHD4, GDE1, GDE7/GDPD3, and GDE4/GDPD1 in adipose tissues of obese mice. We found that Western diet-induced obese mice versus chow-fed lean mice (**Fig. 1B**) had increased *Abhd4* mRNA levels in white (**Fig.1C**) and brown (**Fig. 1D**) adipose tissue. There was no difference in adipose *Nape-pld, Gde1, Gde7/Gdpd3, Gde4/Gdpd1* mRNA levels between lean and obese mice (Data not shown).

### Deletion of ABHD4 promotes adipogenesis *in vitro*

To further examine the functional role of ABHD4 in adipocyte biology, we have generated a ABHD4 KO 3T3-L1 pre-adipocyte. We observed that a small amount of truncated ABHD4 protein was detected by Western blot in 3T3-L1 ABHD4 KO compared to control wildtype (WT) pre-adipocytes (**Fig. 2A**). During adipogenesis, fibroblast-like pre-adipocytes differentiate into lipid-laden and insulin-responsive adipocytes. Adipogenesis is a complex and multi-step process involving activation of a cascade of transcription factors such as CCAAT/enhancer binding proteins (C/EBPs), Peroxisome proliferator-activated receptors (PPARs), and Sterol regulatory element binding proteins (SREBPs) that induce gene expression including Acetyl-CoA carboxylase 1 (ACC1) and Fatty acid synthase (FASN) for *de novo* fatty acid synthesis, Fatty acid-binding protein 4 (FABP4) for fatty acid chaperones, CD36 for fatty acid uptake, as well as Glycerol phosphate acyltransferase (GPAT), Acylglycerolphosphate acyltransferase (AGPAT), and Diacylglycerol acyltransferase (DGAT) for fatty acid esterification to form TAG, all of which lead to adipocyte development(28, 29).

**Fig. 2.**
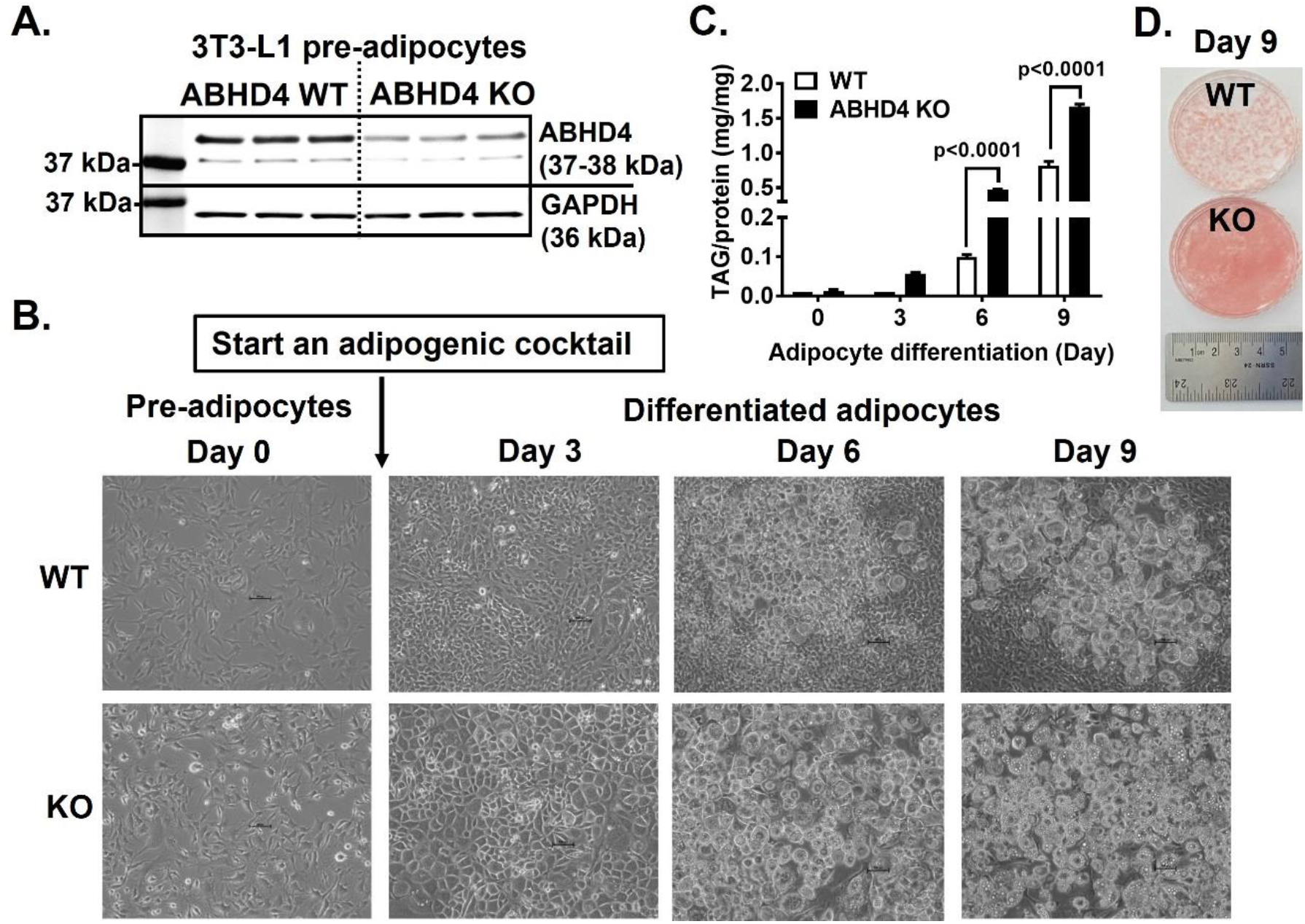
The effect of ABHD4 deletion on adipocyte differentiation. **(A)** Cellular proteins of control wildtype (WT) and ABHD4 knockout (KO) 3T3-L1 pre-adipocytes (n=3/genotype) were harvested and subjected to Western blot using anti-ABHD4 and anti-GAPDH antibodies. **(B)** WT and ABHD4 KO 3T3-L1 cells (n=3/genotype) were proliferated for 2 days (Day 0) and then differentiated into adipocytes for 9 days. Microscopic images at 10X magnification were captured using a phase contrast microscope with a scale of 100 μm. **(C)** Day 0, 3, 6, and 9 cells were lipid-extracted to measure triacylglycerol (TAG) mass by colorimetric assays. Results are mean ± SEM and analyzed using a two-way ANOVA with Sidak multiple comparisons. **(D)** The TAG in Day 9 cells was stained with Oil Red O and photographed to show Oil Red O staining.

We found that during adiponenic stimuli, ABHD4 KO cells acquired adipocyte morphology and accumulate TAG faster than WT cells (**Fig. 2B**); TAG content was 10-fold higher in ABHD4 KO than in WT cells at Day 3, 5-fold higher at Day 6 (p<0.0001), and 2-fold higher at Day 9 (p<0.0001) of adipocyte differentiation (**Fig. 2C**). The TAG in Day 9 adipocytes was stained with Oil Red O (**Fig. 2D**). WT cells showed typical 3T3-L1 cell culture morphology with a cluster of adipocytes showing red color surrounded by non-differentiated, non-stained fibroblast-like cells (**Fig. 2D**). All of ABHD4 KO adipocytes had TAG lipid droplets at Day 9 (**Fig. 2D**). These data suggest that ABHD4 deletion in 3T3-L1 cells increases adipogenesis and lipid accumulation.

We then examined several key genes that play a role in adipogenesis and lipid metabolism. ABHD4 KO versus WT adipocytes has increased mRNA levels of transcription factor *Pparγ*, but not *C/ebp* (Data not shown) and *Srebf* (i.e., SREBP1) family, during adipocyte differentiation (**Fig. 3**). In addition, except *Acc1* and *Agpat1*, the mRNA levels of *Fasn, Fabp4, Cd36, Agpat9* (i.e., GPAT3), *Agpat6* (i.e., GPAT4), *Agpat2, Dgat1*, and *Dgat2* were increased in ABHD4 KO compared to WT adipocytes during Day 3-9 of differentiation (**Fig. 3**).

**Fig. 3.**
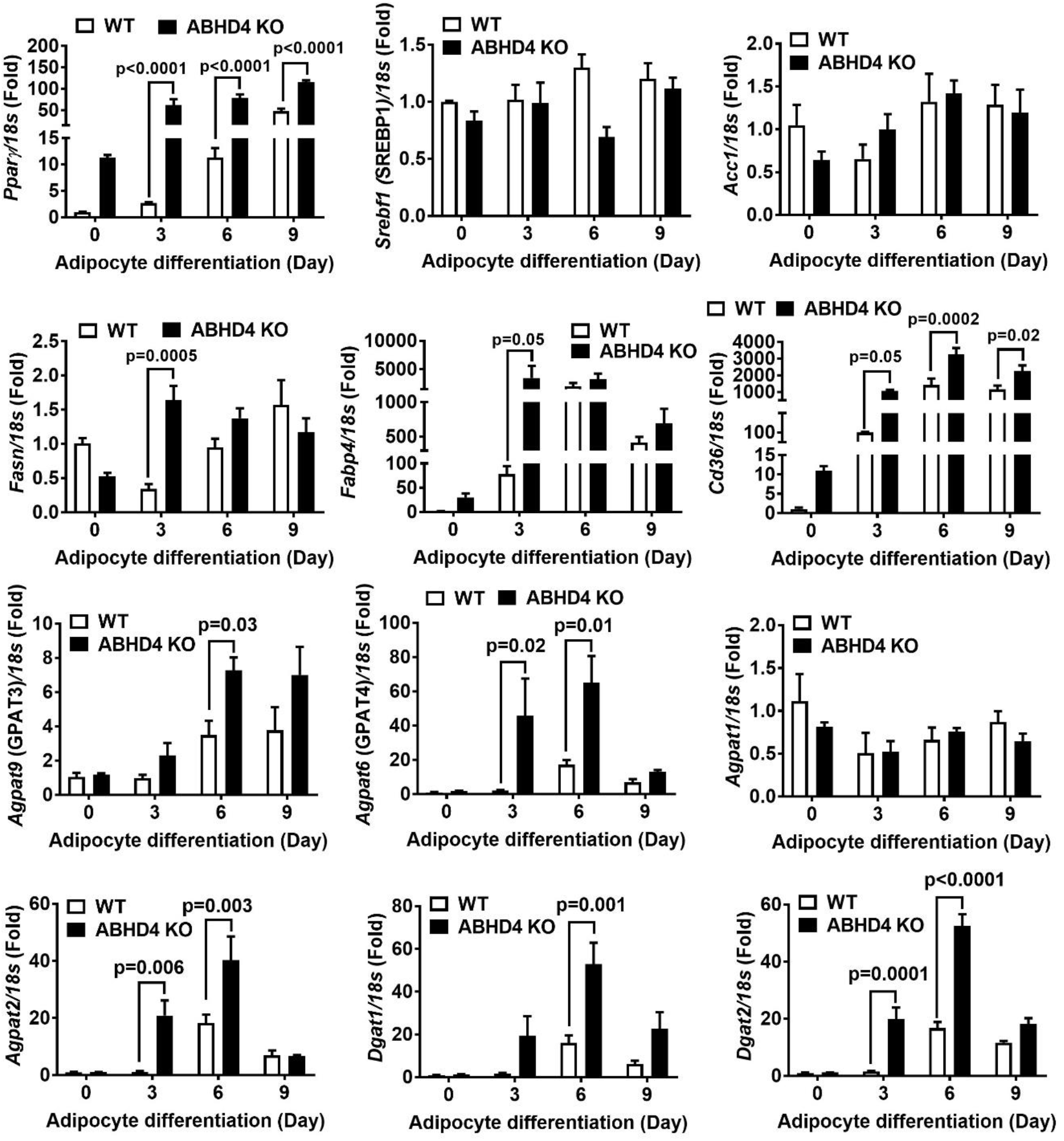
The effect of ABHD4 deletion on adipogenic and lipogenic gene expression. Cellular RNA was extracted from wildtype (WT) and ABHD4 knockout (KO) 3T3-L1 cells (n=3/genotype) and reverse-transcribed into cDNA for real-time PCR quantification of *Pparγ, Srebf1, Acc1, Fasn, Fabp4, Cd36, Agpat9, Agpat6, Agpat1, Agpat2, Dgat1, and Dgat2* normalized to *18s* (endogenous control). Results are mean ± SEM, presented as the fold change compared to WT at Day 0, and analyzed using a two-way ANOVA with Sidak multiple comparisons.

### Deletion of ABHD4 promotes lipogenesis *in vitro*

Next, we performed functional characterization. Day 7 differentiated WT and ABHD4 deficient adipocytes were treated with [^14^C]-acetic acid to determine *de novo* lipogenesis or [^3^H]-oleic acid to measure fatty acid uptake and esterification. Rate of TAG, free cholesterol (FC), cholesteryl ester (CE), and phospholipid (PL) synthesis from [^14^C]-acetic acid was significantly increased (p<0.0001) in ABHD4 deficient adipocytes versus control WT adipocytes (**Fig. 4A**). ABHD4 deficient adipocytes also had significantly increased cellular uptake of [^3^H]-oleic acid (p=0.04) and incorporation of [^3^H]-oleic acid into TAG (p<0.0001), CE (p=0.001), and PL (p=0.0004) compared to control WT adipocytes (**Fig. 4B**). These data demonstrate that ABHD4 deficiency increases lipogenesis, supporting the cell phenotype and gene expression results.

**Fig. 4.**
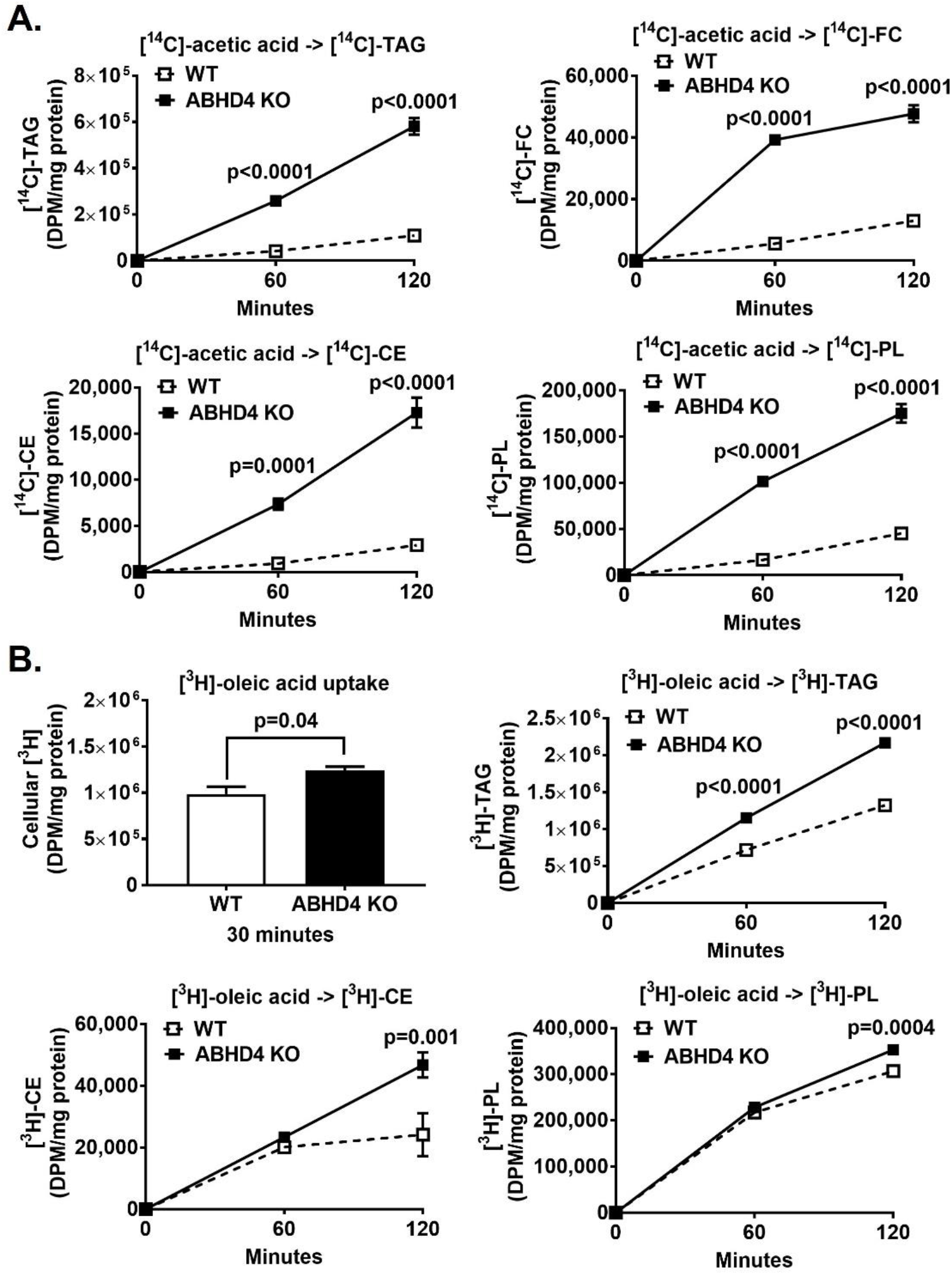
The effect of ABHD4 deletion on lipogenesis. Day 7 differentiated wildtype (WT) and ABHD4 knockout (KO) 3T3-L1 adipocytes (n=3/genotype) were treated with **(A)**0.5 μCi/ml of [1,2-^14^C]-acetic acid or **(B)** 5 μCi/ml of [9,10-^3^H(N)]-oleic acid at the indicated times. Cells were lipid-extracted, and triacylglycerol (TAG), free cholesterol (FC), cholesteryl ester (CE), and phospholipid (PL) were separated using thin layer chromatography. [^14^C]-TAG, [^14^C]-FC, [^14^C]-CE, [^14^C]-PL, cellular [^3^H], [^3^H]-TAG, [^3^H]-CE, and [^3^H]-PL were quantified by liquid scintillation counting. Results are mean ± SEM and analyzed using a two-way ANOVA with Sidak multiple comparisons (i.e., [^14^C]-TAG, [^14^C]-FC, [^14^C]-CE, [^14^C]-PL, [^3^H]-TAG, [^3^H]-CE, and [^3^H]-PL) and a two-tailed Student’s unpaired *t*-test (i.e., cellular [^3^H]).

### Both male and female adipocyte-specific ABHD4 KO mice have a similar metabolic phenotype as their littermate controls on chow or a high-fat diet

We wanted to further investigate whether deletion of ABHD4 promotes adipogenesis and lipogenesis *in vivo* by using adipocyte specific ABHD4 KO (Abhd4^adipose−/−^) mice and their control littermates (Abhd4^adipose+/+^). The genotype of mice was verified by standard PCR and quantitative real-time PCR (**Fig. 5A**) methods. In a similar fashion to data obtained from ABHD4 KO 3T3-L1 (**Fig. 2**), stromal vascular cells isolated from the epidydimal visceral white adipose tissue of chow-fed male Abhd4^adipose−/−^ mice accumulated more TAG (2-fold, p=0.05) than their littermate control male Abhd4^adipose+/+^ mice (**Fig. 5B**). However, there was no difference in body weight and metabolic outcomes such as glucose tolerance test (GTT) between male or female Abhd4^adipose−/−^and Abhd4^adipose+/+^ mice on chow (**Fig. 5C**). We then challenged male mice with a high fat diet. There was no difference in body weight and GTT (**Fig. 5D**) as well as indirect calorimetry measurements (**Fig. 5E**) including oxygen consumption, carbon dioxide production, respiratory exchange ratio, food intake, energy expenditure, locomotor activity, and ambulatory activity (Data not shown) between Abhd4^adipose−/−^ and Abhd4^adipose+/+^ mice on a high fat diet. Please note that despite mice having the C57BL/6 background, well known for being susceptible to diet-induced obesity, we only observed a mild weight gain in our male mice (**Fig. 5D**) and female mice were resistant to diet-induced obesity (Data not shown). Collectively, these data suggest that deletion of ABHD4 in adipocytes does not affect adipose and systemic metabolism in mice fed chow or a high fat diet.

**Fig. 5.**
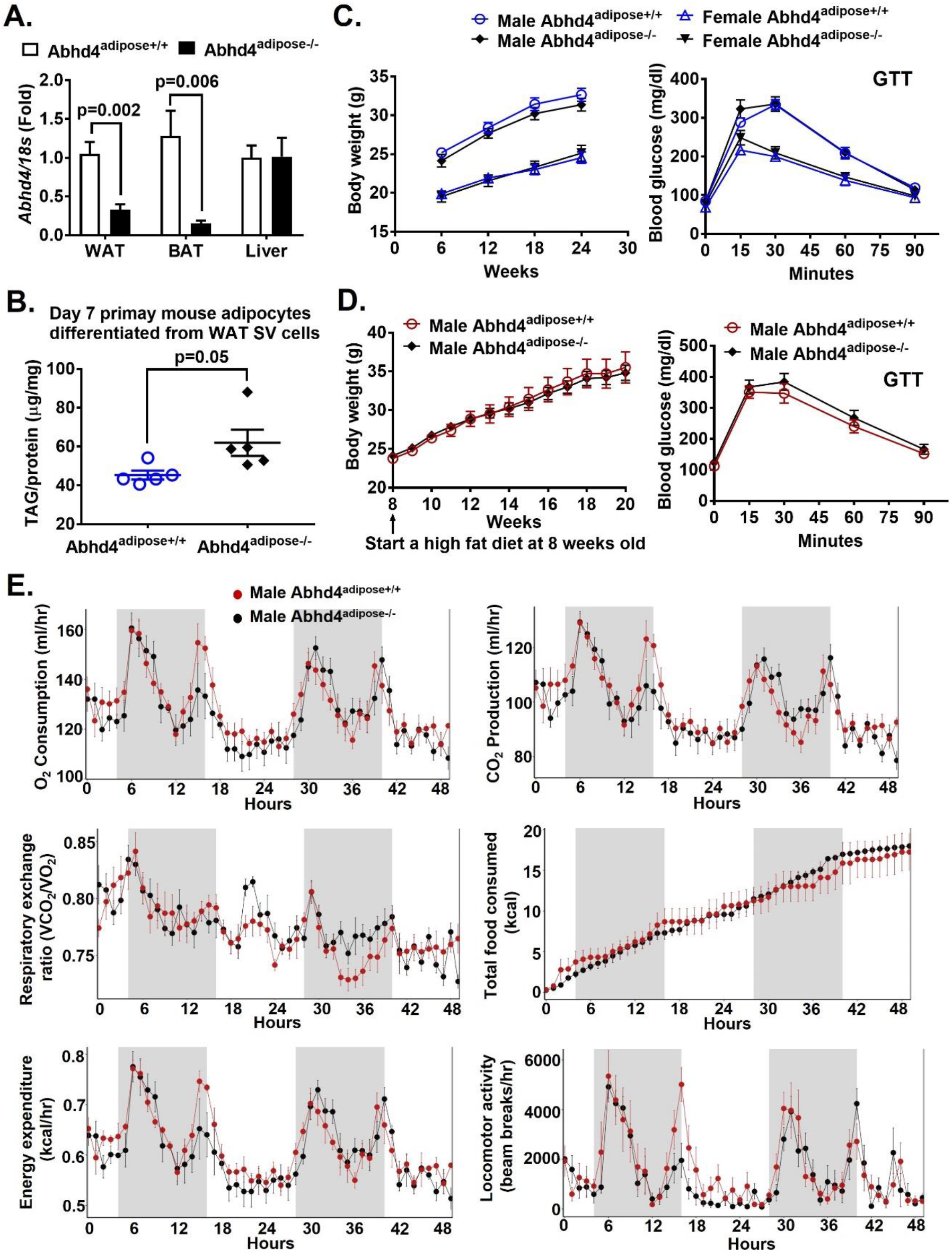
Metabolic phenotype of adipocyte specific ABHD4 knockout (Abhd4^adipose−/−^) versus their control littermate (Abhd4^adipose+/+^) mice. **(A)** Gonadal visceral white adipose tissue (WAT), brown adipose tissue (BAT), and liver were harvested from 8 weeks old male (n=3) and female (n=3) Abhd4^adipose+/+^ and Abhd4^adipose−/−^ mice for RNA extraction, cDNA synthesis, and real-time PCR quantification of *Abdh4* normalized to *18s* (endogenous control). Results are mean ± SEM, presented as the fold change compared to Abhd4^adipose+/+^, and analyzed using a two-tailed Student’s unpaired *t*-test. **(B)** Stromal vascular (SV) cells were isolated from epididymal visceral WAT of chow-fed male mice (n=5/genotype), differentiated into adipocytes for 7 days, and then lipid-extracted to measure triacylglycerol (TAG) mass by colorimetric assays. Results are mean ± SEM and analyzed using a two-tailed Student’s unpaired *t*-test. **(C)** Body weights of chow-fed mice (n=7-9/sex/genotype) were measured weekly and presented every 6 weeks. At 24 weeks of age, chow-fed mice were fasted for 16 hours, and glucose tolerance test (GTT) was conducted. Results are mean ± SEM and analyzed using a two-way ANOVA with Sidak multiple comparison. **(D)** Eight weeks old male mice (n=7-8/genotype) were fed a high fat diet for 12 weeks. Body weights were measured weekly. After a 12-week of a high fat diet feeding, at 20 weeks of age, mice (n=5/genotype) were fasted for 16 hours, and GTT was conducted. Results are mean ± SEM and analyzed using a two-way ANOVA with Sidak multiple comparisons. **(E)** After a 12-week of a high fat diet feeding, at 20 weeks of age, male mice (n=3/genotype) were used for indirect calorimetry (TSE PhenoMaster System). The graphs were created and statistical analysis (one-way ANOVA GLM) was performed using a software package, CalR.

### High fat-fed male and female whole body ABHD4 KO mice have a similar metabolic phenotype as their littermate controls

We then investigated whether ABHD4 deletion in pre-adipocytes or fetal development disrupts adipose and systemic metabolism using a whole body ABHD4 KO (Abhd4^−/−^) mice and their control littermates (Abhd4^+/+^). The genotype of mice including Abhd4^+/+^, Abhd4^+/−^, and Abhd4^−/−^ was verified by standard PCR and quantitative real-time PCR (**Fig. 6A**). Similar to the metabolic phenotypes of adipocyte specific ABHD4 KO mice, we observed no difference in body weight and GTT between male or female Abhd4^−/−^ and Abhd4^+/+^ mice on a high fat diet (**Fig. 6B**). There was also no difference in mouse fat and lean mass quantified by EcoMRI between male or female Abhd4^−/−^ and Abhd4^+/+^ mice (**Fig. 6C**). Furthermore, the individual fat pad mass was unaffected by ABHD4 deficiency (**Fig. 6D**). Interestingly, male Abhd4^−/−^ mice consumed more food (kcal, p=0.009) but also had increased energy expenditure (kcal/hr, p=0.04) compared to their littermate control male Abhd4^+/+^ mice (**Fig. 6E**), thus the body weight/composition did not differ between the two mouse genotypes (**Fig. 6B-D**).

**Fig. 6.**
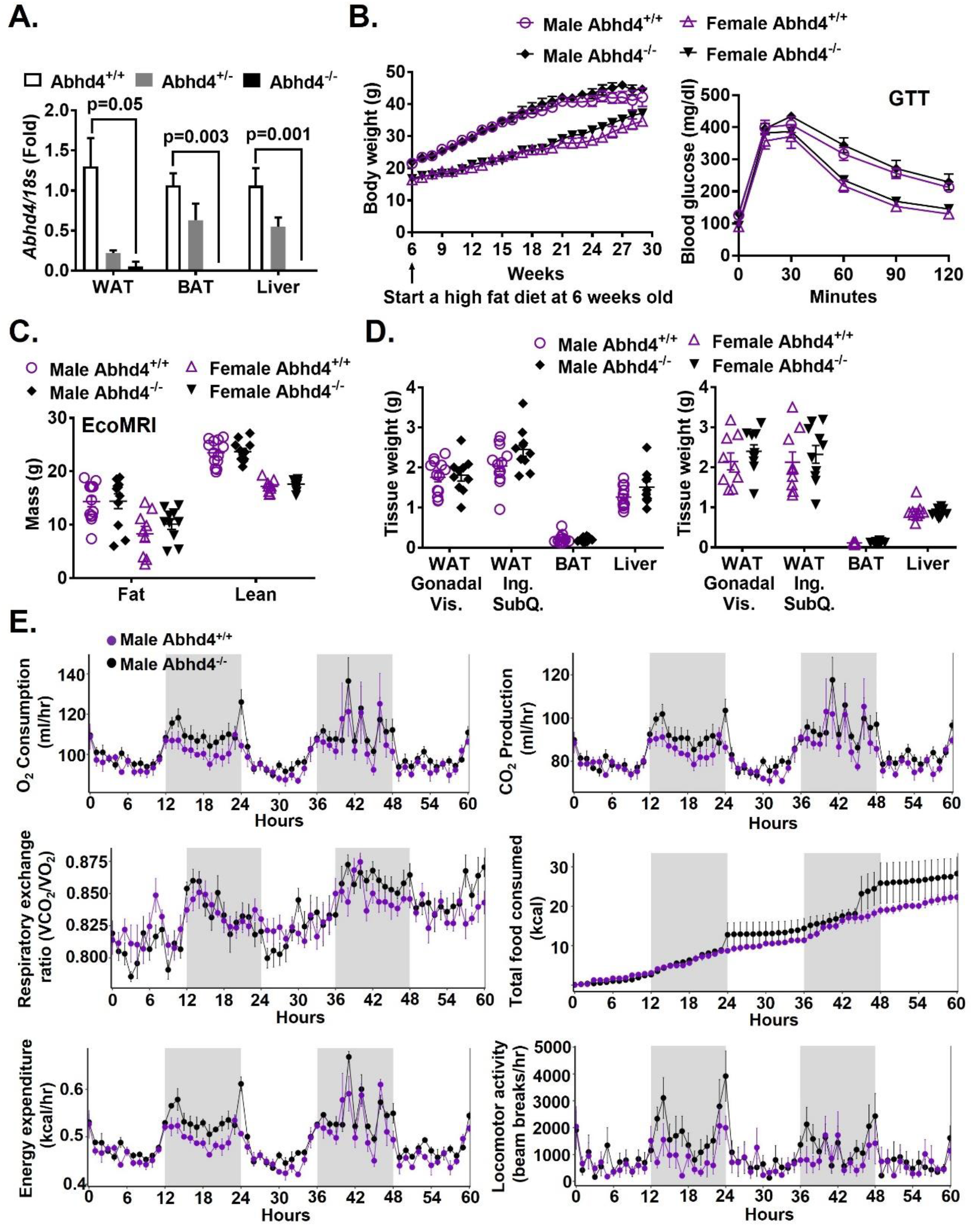
Metabolic phenotype of whole body ABHD4 knockout (Abhd4^−/−^) versus their control littermate (Abhd4^+/+^) mice. **(A)** Epididymal visceral white adipose tissue (WAT), interscapular brown adipose tissue (BAT), and liver were harvested from 8 weeks old male Abhd4^+/+^, Abhd4^+/−^, and Abhd4^−/−^ mice (n=3-7) for RNA extraction, cDNA synthesis, and real-time PCR quantification of *Abdh4* normalized to *18s* (endogenous control). Results are mean ± SEM, presented as the fold change compared to Abhd4^+/+^, and analyzed using a one-way ANOVA with Dunnett’s multiple comparisons. **(B)** Six weeks old male (n=10-13/genotype) and female (n=9-10/genotype) mice were fed a high fat diet for 24 weeks. Body weights were measured weekly. After a 16-week of a high fat diet feeding, at 22 weeks of age, mice were fasted for 16 hours, and glucose tolerance test (GTT) was conducted. Results are mean ± SEM and analyzed using a two-way ANOVA with Sidak multiple comparisons. **(C)** Body composition of mice fed a high fat diet for 15 weeks, at 21 weeks of age, were quantified by EcoMRI. Results are mean ± SEM and analyzed using a two-tailed Student’s unpaired *t*-test. **(D)** After mice were on a high fat diet for 24 weeks, at 30 weeks of age, their gonadal visceral (Vis.) WAT, inguinal (Ing.) subcutaneous (SubQ.) WAT, BAT, and liver were then collected and weighted. Results are mean ± SEM and analyzed using a two-tailed Student’s unpaired *t*-test. **(E)** After a 17-week of a high fat diet feeding, at 23 weeks of age, male mice (n=3/genotype) were used for indirect calorimetry (TSE PhenoMaster System). The graphs were created and statistical analysis (one-way ANOVA GLM) was performed using a software package, CalR.

### Deletion of ABHD4 did not affect NAE production in adipose tissue

Since ABHD4 plays a role in NAE production (**Fig. 1A**), we wanted to investigate whether deletion of ABHD4 in adipocytes affects NAE such as oleoylethanolamide (OEA) levels in adipose tissues of mice. The rationale to study OEA is because Piomelli and colleagues discovered that feeding induces intestinal NAPE-PLD expression and activity, which cleaves NAPE (**Fig. 1A**) such as *N*-oleoyl-phosphatidylethanolamine (NOPE), producing OEA, one of NAE molecules (30, 31). PPARα is activated by OEA, resulting in gut vagal afferent stimulation and satiety(32, 33). They also found that endogenous OEA production exhibits a diurnal cycle in rat white adipose tissue (i.e., higher OEA levels during the daytime or fasting since mice are nocturnal)(34), in contrast to feeding-induced OEA production in the gut, suggesting tissue-specific stimuli may affect OEA production. Furthermore, exogenous OEA treatment of freshly isolated rat adipocytes or *in vivo* treatment of rats and mice with OEA stimulates lipolysis, which does not occur in adipocytes or mice lacking PPARα(35). However, enzymes that control adipose OEA turnover and OEA-mediated lipolysis in response to a diurnal cycle or fasting status have not been characterized.

We first performed a fasting and refeeding study to investigate the association between the gene expression of enzymes in the classical pathway (i.e., *Nape-pld*) versus the alternative pathway (i.e., *Abhd4, Gde1, Gde7/Gdpd3*, and Gde4/*Gdpd1*), and the production as well as function of OEA in adipose tissues. We found that *Abhd4* and *Gde1* (i.e., the alternative pathway), but not *Nape-pld* (i.e., the classical pathway), mRNA levels were increased in epididymal visceral white (**Fig. 7A**) and interscapular brown (**Fig. 7B**) adipose tissue of chow-fed male C57BL/6J mice after a 24-hour fast compared to 24-hour fasting/12-hour refeeding. A similar trend in adipose gene expression was also observed in chow-fed female C57BL/6J mice (**Fig. 7C-D**). The mRNA level of *Gde7/Gdpd3* was relatively low in both adipose tissues (**Fig. 7A-D**). *Pparα* and *Pparγ* gene expression in adipose tissues was known to be regulated by fasting and refeeding, therefore these genes served as controls here (36, 37). We further found that fasting versus refeeding induced OEA production in both white (**Fig. 7E**) and brown (**Fig. 7F**) adipose tissues, and enhanced lipolysis, indicated by elevated circulating free fatty acid concentrations (**Fig. 7G**). These data demonstrate that the alternative pathway is predominant in adipose tissues (i.e., relatively high mRNA levels of *Abhd4* and *Gde1*), and there is a positive association between adipose *Abhd4* expression and OEA production and lipolysis in the fasted state. We then hypothesized that Abhd4^adipose−/−^ mice will have decreased adipose OEA content and lipolysis during fasting. However, we did not observe any difference in fasted adipose OEA levels and serum free fatty acid concentrations in male Abhd4^adipose−/−^ mice compared to Abhd4^adipose−/−^ on chow (data not shown) or fed a high fat diet (**Fig. 7H-J**).

**Fig. 7.**
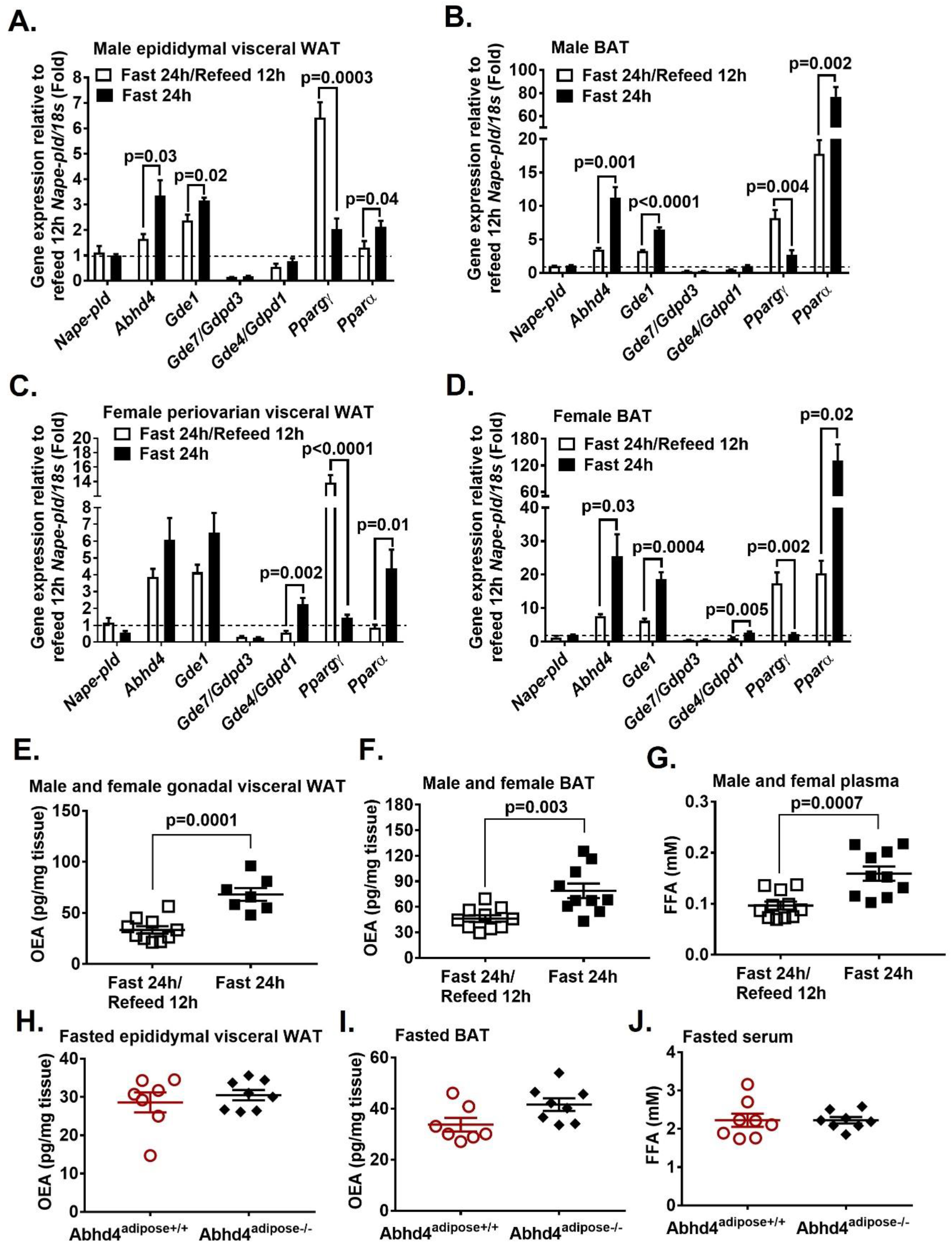
ABHD4’s role in adipose oleoylethanolamide (OEA) production. Sixteen-week-old **(A-B)** male and (**C-D**) female C57BL/6J mice were fasted for 24 hours (n=5/sex) or fasted for 24 hours and then refed chow for 12 hours (n=5/sex). **(A, C)** Gonadal visceral white adipose tissue (WAT) and **(B, D)** interscapular brown adipose tissue (BAT) RNA was extracted and reverse-transcribed into cDNA for real-time PCR quantification of genes. Results were normalized to *18s* (endogenous control) and displayed as fold change relative to *Nape-pld/18s* of 12-hour refeeding. **(E)** Gonadal visceral WAT (n=7-10) and (**F**) BAT (n=10) was used for quantifying OEA content using mass spectrometry. (**G**) Plasma free fatty acid (FFA) concentrations were measured using an enzymatic colorimetric assay. **(H)**Epididymal visceral WAT, **(I)** BAT, and **(J)** blood were collected from a high fat diet-fed male Abhd4^adipose+/+^ (n=7) and Abhd4^adipose−/−^ (n=8) mice after a 16-hour fast for quantifying **(H-I)** adipose OEA content using mass spectrometry and **(J)** serum FFA concentrations using an enzymatic colorimetric assay. All results are mean ± SEM and analyzed using a two-tailed Student’s unpaired *t*-test.

## Discussion

ABHD4 is a (Lyso)-NAPE lipase (**Fig. 1A**). Human ABHD4 is a 342-residue protein (38.8 kDa) encoded by 8 exons located on chromosome 14q11.2. Mouse ABHD4 is a 337-residue protein with 99% amino acid identity with human ABHD4. The Cravatt lab reported that ABHD4 is highly expressed in mouse brain (12). We used the GTEx RNAseq dataset to examine *ABHD4* mRNA profiles across all human tissues and found that adipose tissues demonstrate the second highest *ABHD4* mRNA expression (reproductive organs have the highest *ABHD4* mRNA expression)(38). We further examined the relationship between adipose ABHD4 expression and obesity. Our collaborators observed that ABHD4 gene expression in human subcutaneous adipose tissue (located beneath the skin) is positively correlated with Body Mass Index (BMI, kg/m^2^) in the African American Genetics of Metabolism and Expression (AAGMEx) cohort (r=0.13, p=0.0000365), a cohort of 256 African Americans from North Carolina with in depth glucometabolic phenotyping and adipose tissue transcriptome analysis(39). In addition to the human cohorts, adipose *Abhd4* expression positively correlates with retroperitoneal visceral (located in the abdominal cavity) fat pad weight (n=430, r=0.15, p=0.002) in an outbred rat model(39). The human cohort, an outbred rat model, and our diet-induced obese mouse study (**Fig. 1B-D**) demonstrate for the first time that adipose ABHD4 gene expression is positively associated with obesity, suggesting a previously unappreciated role for ABHD4 in controlling adipose TAG homeostasis.

To better understand the role of ABHD4 in adiposity, we generated an ABHD4 KO mouse 3T3-L1 pre-adipocyte using CRISPR gene editing. Our cell study shows for the first time that deletion of ABHD4 in 3T3-L1 pre-adipocytes promotes adipogenesis and lipogenesis (**Fig. 2–3**). The underlying mechanisms may go beyond simply catabolizing NAPE and synthesizing NAE as we observe no difference in adipose NAE such as OEA production and the overall metabolic phenotypes between adipocyte specific ABHD4 KO mice or whole body ABHD4 KO mice and their littermate control mice on chow or a high fat diet (**Fig. 4–7**). Indeed, the Cravatt lab found that although GP-NAE (the product) decreased, neither NAPE (the substrate) nor NAE (the alternative pathway in **Fig. 1A**) was significantly affected in the brain of whole body ABHD4 KO mice(21). This may be due to the direct conversion from NAPE to NAE by NAPE-PLD (the classical pathway in **Fig. 1**) that controls steady-state concentrations of these lipids.

How does deletion of ABHD4 promote adipogenesis and lipogenesis if NAPE and NAE metabolism is not responsible for this cell phenotype. A recent study surveying protein-protein interactions across the adult mouse brain identified that ABHD4 is physically binding to the regulatory β subunits of Protein kinase CK2 (previously called Casein kinase 2 or CK-II), possibly regulating the protein stability and enzyme activity of CK2(40). Protein kinase CK2 is a ubiquitously expressed and constitutively active serine/threonine protein kinase(41). Structurally, mammalian Protein kinase CK2 is a tetrameric enzyme, composed of two catalytic (α and α’) and two regulatory (β) subunits. The regulatory β subunit does not control catalytic activity of the CK2α or CK2α’ subunits. CK2β regulates CK2’s protein stability(42), substrate specificity(43), and the ability to attach and penetrate the cell membrane(44). The constitutively active Protein kinase CK2 phosphorylates hundreds of protein substrates and controls numerous cellular processes including adipocyte differentiation(41, 45).

Previous studies have demonstrated that CK2 protein expression and kinase activity was induced during the early stage of 3T3-L1 adipogenesis (< Day 6), but after Day 6, alteration of CK2 expression and activity by CK2 inhibitors have no effect on adipocyte maturation(46–48). In addition, the transcription factor C/EBPδ has been identified as a phosphorylated substrate for CK2 to promote early adipogenesis by increasing *Pparγ* mRNA expression(49). Therefore, our future study will test a hypothesis that ABHD4 regulates the early stage of adipocyte differentiation by acting as a suppressor to Protein kinase CK2. We have generated a 3T3-L1 cell line in which FLAG is knocked in to tag endogenous ABHD4 (as confirmed by immunoblotting, unpublished) for this future study. We propose that ABHD4 not only works as a (Lyso)-NAPE lipase, but can also physically interact with other proteins such as Protein kinase CK2 in pre-adipocytes to retard adipocyte differentiation. Overall, in this study, we demonstrate that ABHD4 functions to regulate adipocyte differentiation in vitro, but ABHD4 is dispensable for adipose tissue expansion in high-fat diet fed mice.

## Author contributions

M.S. and S.S. acquisition and analysis of mouse data; A.O.C and C.M.F. acquisition and analysis of mass spectrometry data; C.C.K. writing-original draft, conception and design, acquisition and analysis of cell data, interpretation of data, funding acquisition, and final approval of the version to be published

## Acknowledgements

The authors wish to acknowledge the support of the Wake Forest Baptist Comprehensive Cancer Center Proteomics and Metabolomics Shared Resource, supported by the National Cancer Institute’s Cancer Center Support Grant award number P30CA012197. The content is solely the responsibility of the authors and does not necessarily represent the official views of the National Institutes of Health.

## Funding and additional information

This work was supported by National Institute of Diabetes and Digestive and Kidney to C.C.K. (DK117069). The authors declare that they have no conflicts of interest with the contents of this article.

---

**Supplementary Fig. 1.**
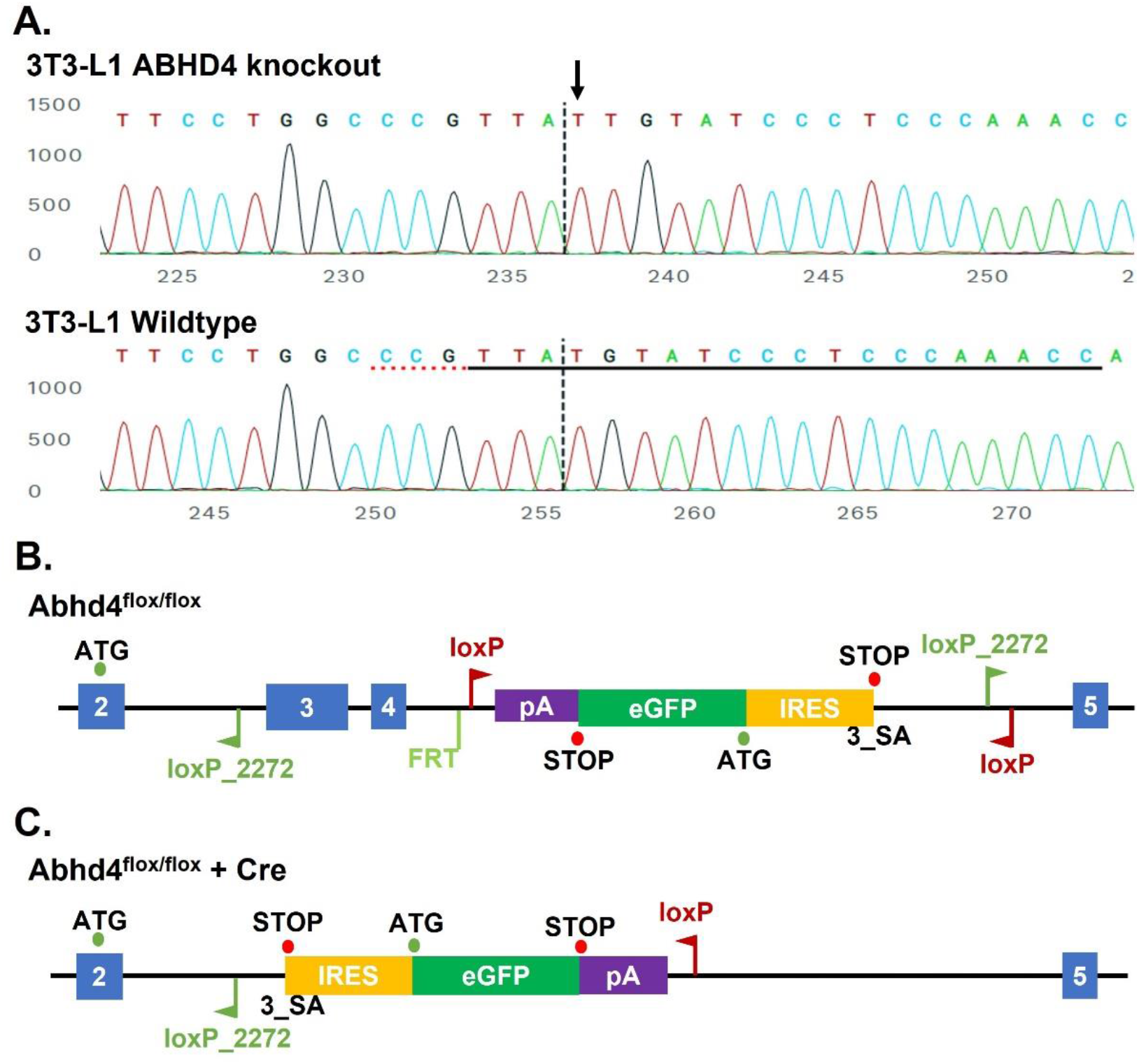
**(A)** The Sanger sequence view showing edited (knockout) and wildtype sequences of Abhd4 in the region around the guide sequence. The horizontal black underlined region represents the guide sequence targeting exon 3 of Abhd4. The horizontal red dotted underline is the PAM site. The vertical black dotted line represents the actual cut site. The knockout clone was cut and had an extra nucleotide insertion (an arrow pointing down) during the non-homologous end joining repair process, resulting in a frameshift mutation that causes premature termination of translation at a new nonsense codon. **(B)** Gene targeted locus of the Abhd4 gene: Exons 3 and 4 (blue boxes) of Abhd4 gene locus and an inverted reporter cassette, 3_SA_IRES_eGFP (3_Splice Acceptor_Internal Ribosome Entry Site_Enhanced Green Fluorescent Protein), were flanking with loxP and loxP_2272 sites via gene targeting in C57BL/6 mouse embryonic stem cells. **(C)** Gene targeted locus after Cre recombinase: Cre recombinase-mediated deletion of the floxed region should excise exons 3-4 and invert the GFP knockin cassette (3_SA_IRES_eGFP) into the correct orientation for expression.

